# Neural systems of cognitive demand avoidance

**DOI:** 10.1101/211375

**Authors:** Ceyda Sayalı, David Badre

**Author notes:** **Address correspondence to:** Ceyda Sayalı Box 1821 Brown University Providence, RI 02912-1978.

## Abstract

Cognitive effort is typically aversive, evident in people’s tendency to avoid cognitively demanding tasks. The ‘cost of control’ hypothesis suggests that engagement of cognitive control systems of the brain makes a task costly and the currency of that cost is a reduction in anticipated rewards. However, prior studies have relied on binary hard versus easy task subtractions to manipulate cognitive effort and so have not tested this hypothesis in “dose-response” fashion. In a sample of 50 participants, we parametrically manipulated the level of effort during fMRI scanning by systematically increasing cognitive control demands during a demand-selection paradigm over six levels. As expected, frontoparietal control network (FPN) activity increased, and reward network activity decreased, as control demands increased across tasks. However, avoidance behavior was not attributable to the change in FPN activity, lending only partial support to the cost of control hypothesis. By contrast, we unexpectedly observed that the deactivation of a task-negative brain network corresponding to the Default Mode Network (DMN) across levels of the cognitive control manipulation predicted the change in avoidance. In summary, we find partial support for the cost of control hypothesis, while highlighting the role of task-negative brain networks in modulating effort avoidance behavior.

## Introduction

Cognitive effort influences our everyday decisions about whether to perform challenging mental tasks. Most people prefer less cognitively effortful tasks when given a choice, a phenomenon known as ‘demand avoidance’ (Kool et al., 2010). However, it is not yet settled what cognitive and neural mechanisms underlie this avoidance behavior, or whether the deployment of these mechanisms varies among individuals.

One account of demand avoidance behavior is the ‘cost of effort’ hypothesis, according to which the brain codes cognitive effort as disutility (Botvinick, 2007). Consistent with this hypothesis, the value of reward is discounted as a function of effort requirements (Westbrook et al., 2013). In the brain, this hypothesis predicts that this cost should be computed by the same networks that typically process reward, such as the mesolimbic dopaminergic system, including the medial prefrontal cortex and the ventral striatum (VS). This prediction is supported by at least one fMRI study which observed reduced activation in VS following effortful task performance (Botvinick et al., 2009). However, the engagement of these systems may be affected by whether one is performing the effortful task or selecting whether to perform it. For example, Schouppe et al. (2014) found that VS increased, rather than decreased, its activity during the selection of the effortful task. Thus, evidence that the brain registers effort as disutility has received limited, albeit conflicting, support.

A second question concerns what makes a task cognitively effortful. An influential hypothesis proposes that tasks are effortful to the degree that they recruit cognitive control. Thus, according to the “cost of control” hypothesis, cognitive control might be recruited to the degree that its associated costs do not exceed its anticipated benefits (Shenhav et al, 2013). Consistent with this hypothesis, people avoid tasks that require more task switching (Kool et al., 2010), greater working memory load (Westbrook et al., 2013), and greater response conflict (Schouppe et al., 2014), all of which putatively require greater cognitive control.

Given the general association of cognitive control with a dorsal fronto-parietal network (FPN) in the brain (Fedorenko et al., 2013; Niendam et al., 2012; Badre & D’Esposito, 2009; Vincent et al., 2008; Cole & Schneider, 2007; Dosenbach et al., 2007), a reasonable prediction is that engagement of FPN by a task will be associated with a tendency to avoid that task. At least one study has observed that participants who tended to avoid demanding tasks also showed increased FPN activity during effortful task performance (McGuire & Botvinick, 2010). However, this study contrasted only two effort levels and so had limited sensitivity to test whether cognitive control demands were associated, in a dose-response fashion, with changes in brain-behavior relationships within-subject. Further, this study focused on univariate activation change in FPN, but other evidence suggests that functional connectivity with the FPN may be important for cognitive effort (e.g., Ray et al., 2017).

Accordingly, in the present study, we sought to provide a more sensitive test of the brain-behavior relationships predicted by the *cost-of-control* hypothesis. Specifically, we developed a parametric version of the Demand Selection Task (DST) paradigm for fMRI to sensitively test how incremental changes in activity and connectivity levels in FPN, the reward network, and/or other systems related to demand avoidance. Further, we addressed how these within-subjects effects were modulated by individual differences in demand avoidance versus demand seeking behavior. Using this approach, we observed only qualified support for the cost for control hypothesis, in that though FPN activity was related to effort avoidance, this relationship was not clearly mediated by the cognitive control manipulation, as opposed to other factors. However, we discovered that effort-related modulation of regions corresponding to the “default mode network” were associated with effort-related changes in effort avoidance.

## Methods

### 1.1. Participants

Twenty-eight right-handed adults (aged 18-35; 14 female) with normal or corrected to-normal vision were recruited for the behavioral demand avoidance experiment (Experiment 1) using Brown research subject pool. Fifty-six right-handed adults (aged 18-35; 26 female) with normal or corrected to-normal vision were recruited for the fMRI experiment (Experiment 2). Two participants withdrew from the fMRI study and did not complete all experimental phases. Two fMRI participants’ data were excluded due to an operator error that resulted in a failure to record behavioral data during scanning. One participant reported during debriefing that they failed to follow instructions. One participant was excluded due to head movement greater than our voxel size across all sessions. So, in total, data from six participants in Experiment 2 were excluded prior to analysis of the fMRI data. Thus, 50 participants were included in the behavioral and fMRI analyses of Experiment 2. All participants were free of neurological or psychiatric conditions, were not taking drugs affecting the central nervous system, and were screened for contraindications for MRI. Participants provided informed consent in accordance with the Research Protections Office at Brown University.

### 1.2. Behavioral Task

In both the Experiment 1 and 2, participants performed a parametric variant of the demand selection task (DST; Fig 1). In an initial Learning phase, the participant associated virtual card “decks”, denoted by particular symbols, with a certain effort level through experience with trials drawn from that deck. Then, in a second Test phase, participants chose which deck they wished to perform (Selection epoch) and then performed the trials drawn from that deck (Execution epoch). A distinguishing characteristic of this version of the DST relative to prior versions is that we varied the effort level parametrically over six levels based on the frequency of task switching required by a deck. This allowed us to characterize the functional form of changes in behavior or fMRI signal due to our cognitive control manipulation. We now provide detail on each of these phases of the behavioral task.

**Figure 1.**
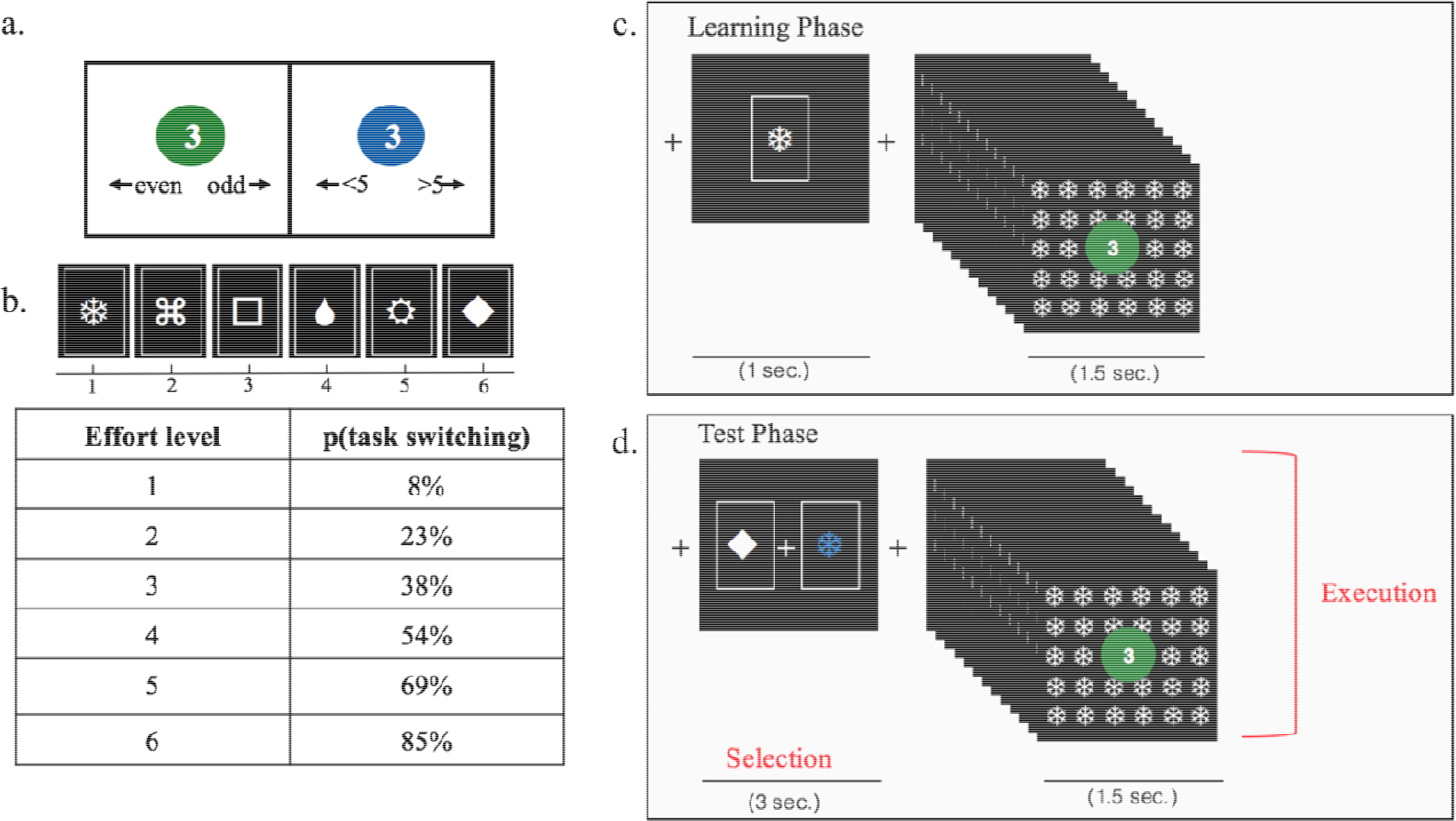
Schematic of the parametric demand selection task. (a) Example trials of the categorization task with a digit printed on a colored circle. The color cued which task to perform. The response mappings shown below the digits are for the reader’s reference and did not actually appear on the screen. (b) An example of six deck cues with their associated effort level pairing, based on the probability of task switching in 13 trials from that deck. (c) Schematic of block events during the Learning Phase. First, a symbol icon is presented for 1 sec, after which the participant completes 13 categorization trials with the associated cue tiled on the background. (d) Schematic of block events during the Test Phase. In the “Selection” epoch, the participant chooses between two symbol icons that are associated with two different effort levels, with a deadline of 3 sec. In the “Execution” epoch, the participant executes the effort level associated with the selected option, while the selected cue is tiled on the background.

#### 1.2.1. The Task “Decks”

Throughout all phases of the experiment, participants performed trials drawn from virtual decks (Fig 1a). Blocks consisted of sequences of 13 consecutive trials drawn from a given deck. On each trial, the participant categorized a presented digit as either odd/even (parity judgment) or greater/less than 5 (magnitude judgment). The color of a circle (green or blue) surrounding the digit cued whether to perform the parity or magnitude judgment on each trial based on a pre-instructed mapping. The participant indicated their categorization by pressing the left or the right arrow key on the keyboard. The response deadline was 1.5 sec. Trials were separated by .2 sec. Response mappings and color-to-task mappings were counterbalanced across participants. Digits 1-4 and 6-9 were used with equal frequency across both tasks and were randomized for order of presentation.

In order to manipulate cognitive effort across decks, we varied the frequency of task switching required by a particular deck (Fig 1b). The probability of switching tasks from trial to trial within a sequence drawn from each deck increased across 6 effort levels: 8%, 23%, 38%, 54%, 69%, 85%. Note that the lowest effort level still included one switch. In this way, all effort levels included the same number of parity and magnitude trials. Thus, any differences in effort preference among these could not be attributed to differences in difficulty or effort between the tasks themselves. Likewise, we did not include “pure blocks” of only parity judgments or only magnitude judgments, as doing so would have made these decks qualitatively different from all other decks, in terms of the amount of each task being performed. Having the lowest effort level include one switch ensures that the only parameter that changes between effort levels is the frequency of task-switching. A higher frequency of task switching is associated with a higher experience of cognitive effort (Monsell, 2003), and has been shown to be aversive (Arrington & Logan, 2004). At the beginning of each block, a shape was presented for 1 sec to indicate which deck was being performed, and this shape was also tiled in the background throughout performance of the block. Participants were told that this shape represented the symbol on the back of the virtual card deck from which trials for that sequence were drawn. Each effort level was associated with a particular deck symbol. Participants were not told about this relationship, but could learn through experience that certain decks were more effortful than others to perform. The mappings between deck symbols and effort levels were randomized across participants.

#### 1.2.2. Practice Phase

During an initial “Practice Phase” participants gained familiarity with the trial structure, the categorization decisions, and the color and response mappings. After being given instructions regarding the categorization and response rules (Fig 1a), they practiced two 13-trial blocks. Each task sequence included the presentation of a deck symbol, and the subsequent performance of 13 consecutive task-switching trials. In the first block of the Practice Phase, feedback was presented after the button press as either ‘Correct’ in pink, ‘Incorrect’ in yellow, or ‘Please respond faster’ if the participant failed to answer in 1.5s. In the second block of the Practice phase, feedback was omitted as would be the case in the actual experiment. The deck symbols that participants encountered in the Practice Phase were for practice only and were not presented during the learning or test phases to avoid any association of error feedback with a particular deck symbol.

#### 1.2.3. Learning Phase

In the Learning Phase (Fig. 1c), participants learned the association between 6 deck symbols and an effort level. Each deck was performed 15 times in random order. In both experiments, this phase was performed in a behavioral testing room outside the magnet.

#### 1.2.4. Test Phase

In the Test Phase (Fig 1d), two decks were presented and the participant chose which they would like to perform (Selection epoch). The participants were told to freely choose between decks prior to a 3 sec deadline. We note that in contrast to other DST variants (Gold et al., 2014), participants in this task were not told about the difficulty manipulation to avoid biasing choices based on participants’ explicit beliefs about effort. Once the participant made their selection, the selected deck turned blue and both options remained on the screen until the end of the 3 sec deadline. In the event of a non-response, the same choice pair was re-presented at the end of the entire experiment until they made a decision. Each pair was selected from the set of fifteen unique (un-ordered) pair combinations of all six decks, excluding self-pairing (e.g., deck #1 paired with deck #1). Each deck was presented either on the left or the right side of the screen, counterbalanced for location across trials. The Selection epoch was followed by the execution of the selected effort level task sequence (Execution epoch). The sequence of events in this epoch was the same as during the Learning phase. We note that based on the self-selected execution of effort tasks in this phase, there were differences in the number of trials each individual performed at each effort level. Although this variability naturally alters our detection power for corresponding effort trials across individuals in the Test Phase, Learning phase ensured that people established stable behavior and effort associations with each level based on the same number of trials with each effort level prior to the Test phase

In the fMRI study, only the Test phase was performed in the MRI scanner, and participants used a MRI-compatible button box to indicate their decisions. The Execution trials were optimized as blocks. The behavioral and fMRI versions of the Test Phase was identical except the Selection and Execution events were separated in time by a jittered time interval (mean 2 secs) so that signal related to each could be analyzed independently. Despite the pseudorandomization procedure adopted for the fMRI study, the mean time intervals (mean 2 secs) between stimuli was identical across behavioral and fMRI versions of the Test Phase. The Test phase was separated into four, approximately 15 minute-long scanning runs. In each run, each pair was presented 3 times in a pseudo-randomly intermixed order, making a total of 180 decision trials across 4 blocks.

#### 1.2.5. Post-experimental Debriefing Inventory

Upon the completion of the experiment, participants were asked to respond to 6 questions regarding their understanding of the task. The first 5 questions required text entry. The final question required participants to enter a number. The questions were as follows: 1) What do you think the deck symbols stood for?, 2) Was there any reason for why you selected certain decks in the last phase?, 3) Did you have a preference for any deck?, 4) What do you think the experimental manipulation was?, 5) Imagine somebody else is about to perform the same experiment. What advice would you give them? And, did you follow this advice?, 6) Please rate each deck symbol in terms of its difficulty (out of 6).

### 1.3. Behavioral Data Analysis

Trials with response times below 200 ms were excluded from further analysis. Execution trials on which participants missed the response deadline were also excluded from further analysis (approximately %1 of execution trials in both phases). Response times were calculated using only correct trials.

Choice behavior was assessed by calculating the probability of selecting an effort level across all selection trials on which that effort level was presented as an option during the Test Phase. The decision difference analyses included the calculation of the choice probability and the decision time to select the easier task across all decisions with the same difference in difficulty levels between effort options in the selection epoch of the Test Phase. For example, choice probability associated with a difficulty difference of 1 would be computed by averaging the probability of choosing the easier task across all choice pairs that differed by 1 effort level (i.e., 1 vs 2, 2 vs 3, 3 vs 4, 4 vs 5 and 5 vs 6).

In order to appropriately test for changes in behavioral selection rates across effort levels while also defining group membership (i.e., Demand Avoiders or Demand Seekers) based on the same choice behavior, we conducted a permutation procedure (see Supplementary Methods 1).

Data were analyzed using a mixed-design analysis of variance (ANOVA) (within subject factor: Effort, between subject factor: Avoidance group). If the sphericity assumption was violated, Greenhouse-Geisser correction was used. Significant interactions were followed by simple effects analysis, the results of which are presented with False Detection Rate (FDR) correction. Alpha level of .05 was used for all analyses. Error bars in all figs stand for within-subject error.

### 1.4. MRI procedure

In the fMRI experiment (Experiment 2), whole-brain imaging was performed with a Siemens 3T Prisma MRI system using a 64-channel head coil. A high-resolution T1-weighted 3D multi-echo MPRAGE image was collected from each participant for anatomical visualization. Each of the four runs of the experimental task involved around between 450 and 660 functional volumes depending on the participant’s response time, with a fat-saturated gradient-echo echo-planar sequence (TR = 2s, TE=28ms, flip angle = 90°, 38 interleaved axial slices, 192 mm FOV with voxel size of 3×3×3 mm). Head motion was restricted with padding, visual stimuli were rear projected and viewed with a mirror attached to the head coil.

### 1.5. fMRI Analysis

Functional images were preprocessed in SPM12 (http://www.fil.ion.ucl.ac.uk/spm). Before preprocessing, data were inspected for artifacts and variance in global signal (tsdiffana, art_global,art_movie). Functional data were corrected for differences in slice acquisition timing by resampling slices to match the first slice. Next, functional data were realigned (corrected for motion) using B-spline interpolation and referenced to the mean functional image. 1-2 sessions were excluded from 3 other participants prior to behavioral analysis due to movement during data collection in the scanner. Functional and structural images were normalized to Montreal Neurological Institute (MNI) stereotaxic space using affine regularization followed by a nonlinear transformation based on a cosine basis set, and then resampled into 2x2x2 mm voxels using trilinear interpolation. Lastly, images were spatially smoothed with an 8 mm full-width at half-maximum isotropic Gaussian kernel.

A temporal high-pass filter of 128 (.0078 Hz) was applied to our functional data in order to remove noise. Changes in MR signal were modeled under assumptions of the general linear model (GLM). Two GLMs were devised: a linear effort-level GLM and an independent effort-level GLM. Both GLMs included nuisance regressors for the six motion parameters (x,y,z,pitch,roll,yaw) and four run regressors for the ‘Linear Effort-Level GLM’ and one run regressor for the ‘Independent Effort Level GLM’. The number of run regressors was different across GLMs because ‘Independent Effort GLM’ included the regressor for each effort level separately. In this GLM, it was not possible to have 4 runs for each effort level since some participants did not choose to select from an effort level an entire run. Thus, in order to retain the entirety of the dataset, we collapsed all trials across runs for each effort level.

#### 1.5.1. Linear Effort-Level GLM

The linear effort-level GLM tested which voxels in the brain parametrically increased or decreased linearly with effort level. Two event regressors were used. Execution events were modeled as a boxcar that onset with the presentation of the first trial stimulus of the sequence and ended with the participant’s response to the final item. Thus, the duration of this boxcar varied as a function of response time. Second, the Selection event regressor modeled each selection choice event with a fixed boxcar of three secs. We used parametric modulators on these event regressors to test the linear effect of effort level. The Execution event regressor was modulated by an Effort Level parametric regressor corresponding to the effort level of that task sequence (1 through 6). The Selection event regressor was modulated by (a) an Effort Level parametric regressor based on the chosen effort level (1 through 6), and (b) a Difference regressor that scaled with the difference between the chosen and the unchosen effort option on that selection trial (1 through 5). Note that as implemented by SPM, each parametric regressor includes only the variance unique to it and shared with those ordered after it. Thus, for example, the Effort Level regressor includes variance explained over and above that shared with the Execution event regressor. The Difference regressor did not yield statistically reliable results and is not discussed further. The Execution and Selection event regressors, along with their parametric modulators, were modeled separately for each scanning run within the GLM.

#### 1.5.2. Independent Effort Level GLM

The independent effort level GLM sought to characterize the signal change related to each effort level independently of each other or of any particular function (e.g., linear). This GLM included twelve event regressors, one for each effort level (1 through 6) by epoch (Execution and Selection). Events in the Execution regressors were modeled as boxcars that onset with the presentation of the first trial stimulus of the sequence and ended with the participant’s response to the final item. Events in the Selection regressors were modeled with an 3 sec boxcar at the onset of each choice pair. The Selection onset regressor was further modulated by a parametric Difference regressor that scaled with the difference between the chosen and the unchosen effort option on that selection trial. Boxcars and durations were the same as in the linear effort level model. In this GLM, four run epochs and a linear drift over the whole experiment were included as nuisance regressors.

For both GLMs, SPM-generated regressors were created by convolving onset boxcars and parametric functions with the canonical hemodynamic response (HRF) function and the temporal derivative of the HRF. Beta weights for each regressor were estimated in a first-level, subject-specific fixed-effects model. For group analysis, the subject-specific beta estimates were analyzed with subject treated as a random effect. At each voxel, a one-sample t-test against a contrast value of zero gave us our estimate of statistical reliability. For whole brain analysis, we corrected for multiple comparison using cluster correction, with a cluster forming threshold of α = .001 and an extent threshold, k, calculated with SPM to set a family-wise error cluster level corrected threshold of p<.05 for each contrast and group (See Table S1). Note that the higher cluster forming threshold helps avoid violation of parametric assumptions such that the rate of false positive is appropriate (Eklund et al., 2016; Flandin & Friston, 2016).

#### 1.5.3. ROI analysis

ROI definition is described below. For each ROI, a mean time course was extracted using the MarsBar toolbox (http://marsbar.sourceforge.net/). The GLM design was estimated against this mean time series, yielding parameter estimates (beta weights) for the entire ROI for each regressor in the design matrix.

##### 1.5.3.1. Independent ROIs

In order to test the cost of control hypothesis, we defined a fronto-parietal control network ROI defined from a functionally neutral group ([Network 12] Yeo et al., 2011) along with a VS ROI (Badre et al., 2014), as VS activity has been consistently observed in cognitive effort literature (Botvinick et al., 2009; Schouppe et al., 2014). We also included a Default Mode Network ROI from a functionally neutral group ([Network 16] Yeo et al., 2011). A priori DMN regions included ventromedial prefrontal cortex (vmPFC), orbitofrontal cortex, posterior cingulate cortex and parts of precuneus. A priori FPN regions included bilateral lateral prefrontal cortex, bilateral parietal cortex and SMA.

##### 1.5.3.2. PCA ROIs

Our study is the first to parametrically manipulate implicitly learned effort cost/values in DST in fMRI. Though we *a priori* conceived of these effort levels as being linear, and so showing a linear brain response, we acknowledge that this need not be the case. As such, we wanted to be sure that our *a priori* expectation of a linear response function with increasing effort, as tested in the parametric GLM, did not cause us to systematically miss networks of the brain that code effort according to a different functional form, but that are relevant signals for the behavioral decision. In order to have an unbiased analysis of the neural response function across effort levels, we adopted a data-driven Principal Component Analysis (PCA) approach on the whole brain (126,866 voxels) to explore the shape of the neural functions of different brain areas that respond to effort execution, and their relation to demand avoidance behavior. We ran the PCA over averaged β estimates of each voxel in the whole brain as they were derived from the Independent Effort Level GLM task Execution Level 1-6 onset regressors across all participants. Therefore, PCA was conducted on a total of 126,866 observations and 6 variables.

We ranked all voxels as a function of PC weight and extracted the first 3000 voxels that loaded the highest (Positive PC) and the least (Negative PC) on the selected components as separate ROIs (Decot et al., 2017). These ROIs further have been analyzed to yield beta estimates for each Positive and Negative PC. The number of voxels to select (3000) was an arbitrary cut off and was not chosen after looking at the results. However, to ensure that outcomes did not depend on this choice, we also tried different numbers of voxels, such as 500 and 10,000, and did not find a qualitative change in average beta estimates ROIs yield.

##### 1.5.3.3. Functional connectivity analysis

For each participant, functional connectivity analysis was implemented in MATLAB using the CONN toolbox (http://www.nitrc.org/projects/conn; <u>Whitfield-Gabrieli & Nieto-Castanon, 2012</u>). CONN uses the CompCor method (<u>Behzadi et al., 2007</u>) which segments white matter (WM) and cerebrospinal fluid (CSF), as well as the realignment parameters entered as confounds in the first-level GLM analysis (<u>Behzadi et al., 2007</u>), and the data were band-pass filtered to 0.01 Hz to 0.1 Hz.

We conducted an ROI-to-ROI analysis to test the functional connectivity between and within the first principal component clusters identified in the PC analysis for the entire sample. PC clusters further have been separated into smaller spatially independent clusters in order to conduct within-PC connectivity analysis. Positive PC1 yielded 4, and Negative PC1 yielded 8 spatially independent clusters. Accordingly, 4 Positive PC1 clusters and 8 Negative PC1 clusters were included as ROIs. Fisher’s Z-transformed correlations were computed for each ROI pair.

Next, we focused on two connectivity measures. First, we calculated the average connectivity Between Positive-Negative PC1, within Positive PC1 and within Negative PC1 clusters at each effort execution level, in order to observe the connectivity change in these measures with increasing effort execution. Next, we collapsed connectivity coefficients for Between Positive-Negative PC1, within Positive PC1 and within Negative PC1 across effort levels and compared average connectivity rates between demand groups. The results of the connectivity analysis did not yield statistically reliable results and they are reported in Supplementary Results 6.

### 1.6. Brain − behavior analysis

In order to understand which brain regions that underlie effort execution predict effortful task selection, all PCs as well as the connectivity measures were related to the selection behavior during the Selection Epoch. We analyzed this relationship in two ways: 1) correlating the change in effort selection rates to the change of brain activity during task execution across effort levels (‘Effort level’), 2) correlating individual effort selection rate to the brain activity during task execution at that effort level (’Individual task selection’), unranked with respect to task switching frequency. The former analysis focuses on activity change in a brain region due to the task-switching manipulation. The latter considers how overall activity levels, independent of a linear task-switching manipulation, correlate with effort avoidance.

For the ‘Effort level’ analysis, we computed the slope of the linear function for the selection rates, β estimates of a PC ROI across effort levels and connectivity coefficients of within and between PCs. We multiplied the slope we obtained from the selection rates with −1, in order to indicate that decreasing likelihood to select more effortful tasks are conceptualized as ‘demand avoidance’ in our task. Then, we correlated the demand avoidance slopes and the β estimate slopes for all Demand Avoiders, in order to address the role of this region in explaining individual variability in demand avoidance behavior.

For the ‘Individual task selection’ analysis, we correlated the β estimate of an ROI at each effort level execution with the selection rate of the corresponding effort level to derive a β value for each participant. This resulting β value indicates the relationship between selecting an effort level given the estimated activity in this region, without presupposing a linear effort manipulation.

In order to test the effects of brain and performance on selection behavior, while accounting for individual variability within demand groups, we adopted the following mixed effects hierarchical regression model:

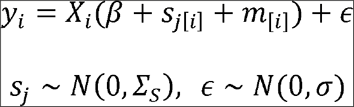

Here, *y*_*i*_ is the *i*th observed selection rate, *X*_*i*_ is a matrix of predictors (brain activity) and covariates (performance), *β*_*i*_ is a vector of coefficients (as in conventional linear regression), *j*[*i*] is the subject of the *i*th observation, so that *S*_*j*[*i*]_ is a subject-specific perturbation of all of the coefficients.

This fitting procedure was implemented using the R package blme (Chung et al., 2013), which includes the lme4 package (Bates et al., 2015) and performs maximum-a-posteriori estimation of linear mixed-effects models.

## 2. Results

### 3.1. *Demand avoidance behavior*

Experiment 1 established the basic behavioral effects of our parametric effort manipulation on performance. Overall, participants were highly accurate on the categorization task across both phases (mean error: 12% in the Learning Phase, 15% in the Test Phase), and participants missed the deadline on few trials (2.3% of Learning phase trials, *SE*=0.003, 1.4% of Test phase trials, *SE*=0.01).

Both RT and error rates showed a linear effect of effort condition, increasing with a higher proportion of task switches across both switch and repeat trials. There was a significant effect of Effort on error rates, (*F*(2.76,74.53)=29.88, *p*< .001, η_p_^2^ =.53) and RT (*F*(2.69,72.71)=131.17, *p*< .001, η_p_^2^ =.83). And, the increase in both was linear over effort levels (errors: *F*(1,27)=61.17, *p*< .001, η_p_^2^ =.69; RT: *F*(1,27)=248.19, *p*< .001, η_p_^2^ =.90). These increases were also significant for both switch and repeat trials tested separately (switch errors: *F*(1,27)=29.73, *p*< .001, η_p_^2^ =.52; repeat errors: *F*(1,27)=42.59, *p*< .001, η_p_^2^ =.61; switch RT: *F*(1,27)=0.37, *p* < .001, η_p_^2^ =.61; repeat RT: *F*(1,27)=42.89, *p*< .001, η_p_^2^ =.61).

During the Selection phase, 25 of the 28 participants (89%) selected the easier task more than 50% of the time overall (Fig 2A). Further, there was a significant effect of effort level on choice behavior, (*F*(5,135)=7.23, *p*< .001, η_p_^2^ =.21), such that higher effort levels were associated with higher avoidance rates than lower effort levels (Fig 3A). This pattern across effort levels was fit by a linear effect (*F*(1,27)=24.54, *p*<.001, η_p_^2^ =.48). The probability of selecting the easier task also significantly changed depending on the difference between effort levels given as options (*F*(3,81.09)=6.07, *p*=.001, η_p_^2^ =.18), such that larger differences increased the probability of choosing the easier task. This effect was linear across effort levels (*F*(1,27)=16.83, *p*<.001, η_p_^2^ =.38). Decision time (Fig 3A) was mostly unaffected by effort level with only a marginal omnibus effect of effort level on decision time (*F*(2.98,80.56)=2.62, *p*=.06, η_p_^2^ =.09) and no effect of difficulty difference.

**Figure 2.**
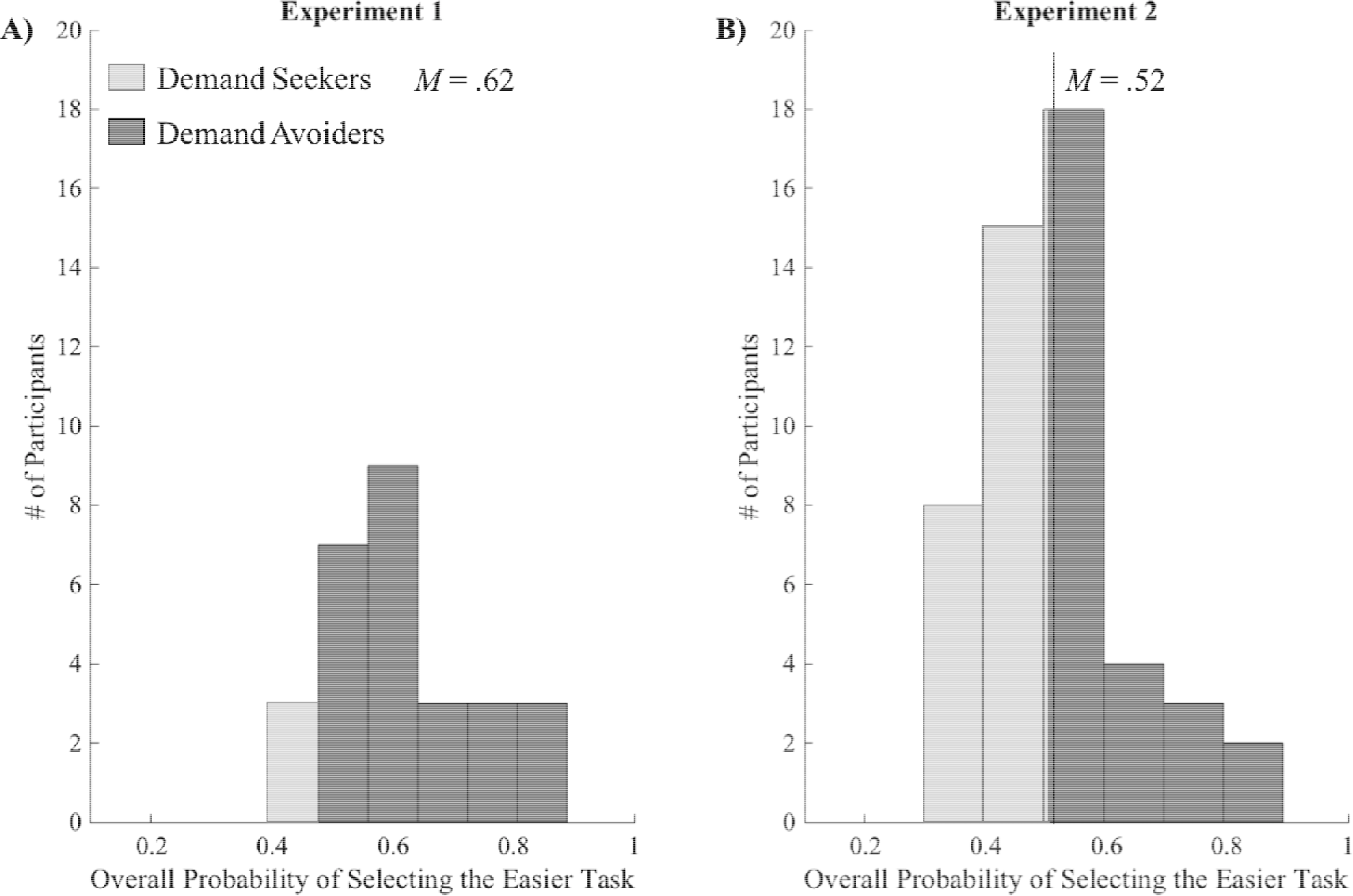
Distribution of participants for the overall probability of selecting the easier task across all decision trials in A) Experiment 1 and B) Experiment 2. In Experiment 1, 25 of the 28 participants (89%) selected the easier task more than 50% of the time overall. In Experiment 2, nearly half of participants (N=24) selected the easier task more than 50% of the time. The mean overall probability of selecting the easier task for each experiment is indicated with a vertical dashed line.

**Figure 3.**
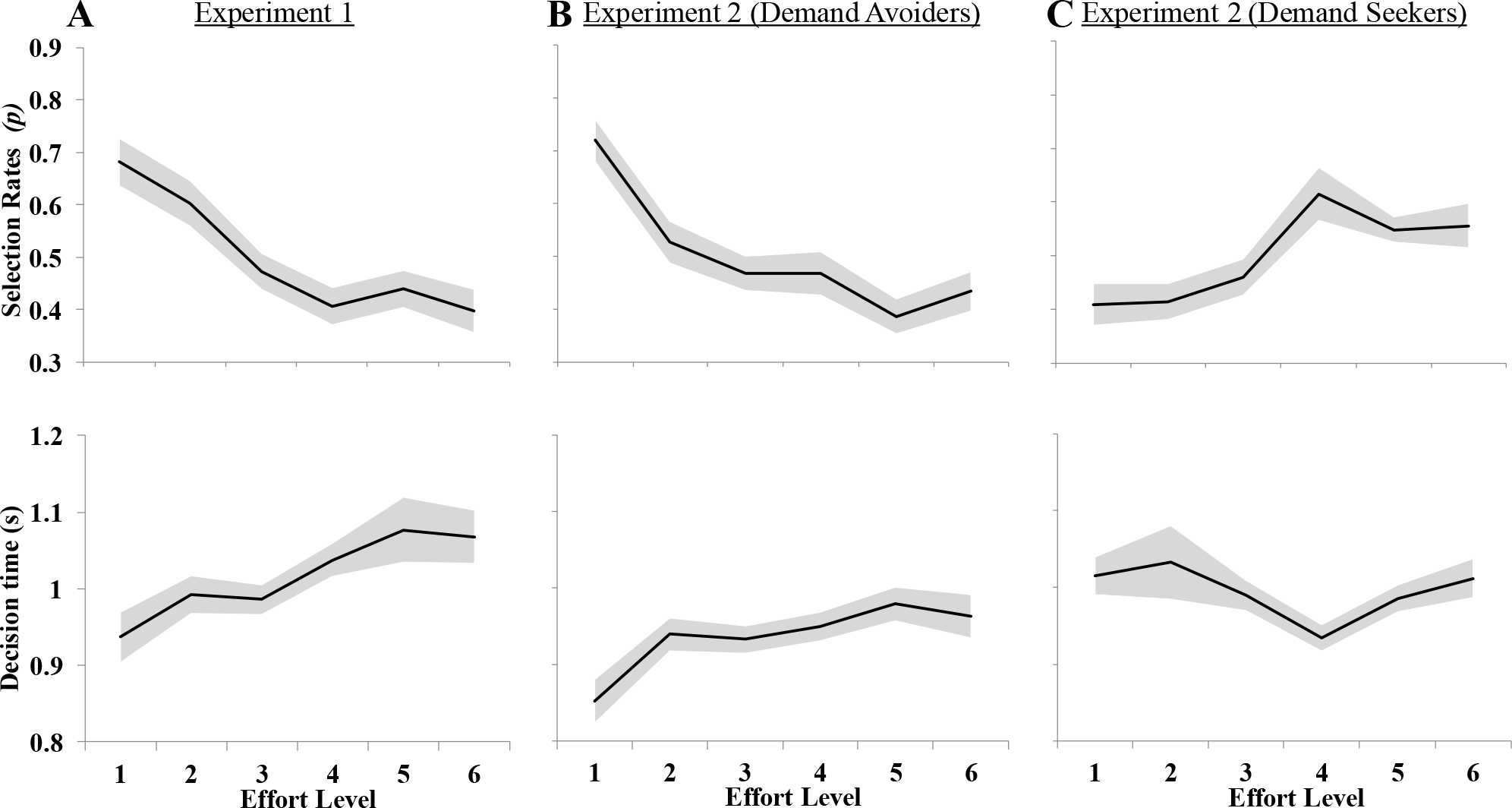
Choice behavior for the entire Experiment 1 sample (A), Demand Avoiders in Experiment 2 (B), and Demand Seekers in Experiment 2 (C). The top panels plot the selection rates across effort levels in terms of the probability of choosing that task when it was given as an option. In Experiment 1, the entire sample showed a decreasing tendency to choose higher effort levels. In Experiment 2, Demand Avoiders showed a decreasing tendency to choose a task as more cognitive effort is required, and Demand Seekers exhibited the opposite pattern. The bottom panels plot the decision times across effort levels. In Experiment 1, decision time was mostly unaffected by effort level with only a marginal omnibus effect of effort level on decision time. In Experiment 2, the decision time to select an effort level linearly increased with increasing effort levels for Demand Avoiders and the decision time to choose an effort level was similar across effort levels for Demand Seekers. All positive and negative trends were significant linear effects at *p* < .05. The shaded area plots standard error of the mean.

Experiment 2 applied the same behavioral task as in Experiment 1, but the Test phase was scanned using fMRI. As with Experiment 1, performance on the categorization task was strong overall (mean error: 12%) with relatively few omissions (<1.5% of trials). Also as with Experiment 1, both RT and error rates showed a linear effect of effort, indicating that our effort manipulation was successful in increasing task difficulty (errors: *F*(1,49)=49.11, *p*< .001, η_p_^2^ =.50; RT: *F*(1,49)=190.48, *p*< .001, η_p_^2^ =.80). And, these effects were evident on both Switch and Repeat trials except for Switch RT (switch errors: *F*(1,49)=47.67, *p*< .001, η_p_^2^ =.49; repeat errors: *F*(1,49)=20.37, *p*< .001, η_p_^2^ =.29; switch RT: *F*(1,49)=0.36, *p* = .55, η_p_^2^ =.01; repeat RT: *F*(1,49)=635.24, *p*< .001, η_p_^2^ =.93)

Unlike in Experiment 1, Experiment 2 participants did not consistently avoid the effortful task as a group during the Selection phase. Rather, nearly half of participants (N=24) selected the harder task more than 50% of the time (Fig 2B). This suggests that, as a group, participants were not avoiding the effortful change more than one expects by chance.

Importantly, however, the diagnostic feature of demand avoidance in our paradigm is the linear change in selection rates across effort levels, as observed in Experiment 1. Therefore, if the subgroup with overall easier task selection rates above 50% (Demand Avoiders) are engaged in effort avoidance, we should also see a linear change in their selection rates across effort levels as was observed in Experiment 1. Likewise, those with selection rates below 50% (Demand Seekers) might show the opposite pattern, with a linearly increasing tendency to choose the harder task. Using a permutation procedure and independent samples of the data (see Supplementary Methods 1 for details), we showed that these selection rate patterns were consistent with participants’ demand group and were not explainable in terms of chance or a selection bias (see Supplementary Results 1). Rather Demand Avoidance versus Demand Seeking appeared to reflect an individual difference.

Our analysis showed that overall demographics between demand groups was similar. There were no age (Demand Avoiders: *M*= 22.12, *SD*= 3.20; Demand Seekers: *M*= 21.45, *SD*= 2.99) or gender differences (Demand Avoiders: N_female_=11, N_male_=15; Demand Seekers: N_female_=14, N_male_=10) between demand groups (age: *F*(1,49)=0.56, *p*= .46; gender: X^2^ (1, N = 50) = 1.28, p=40).

Post-experimental debriefing inventory results showed that participants were mostly unaware of the difficulty manipulation across effort tasks. Only 11 out of 50 participants reported that they noticed a difficulty difference between decks. Further, free text responses showed that even the “aware” participants were not completely sure about the exact difficulty ranking across decks. The final inventory asked participants to rate the difficulty of each deck out of 6. Across participants and decks, average rating accuracy was 15%, consistent with chance level or 1 out of 6 rated correctly.

RTs but not error rates (Table S2) across effort levels and experimental phases differed across demand groups (errors: *F*(1,48)=1.81, *p*= .19, η_p_^2^ =.04; RT: *F*(1,48)=6.21, *p*= .02, η_p_^2^ =.12). Demand Avoiders had faster RTs compared to Demand Seekers in both phases. Both RT and error rates showed a linear effect of effort, indicating that our effort manipulation was successful in increasing task difficulty (errors: *F*(1,48)=72.43, *p*< .001, η_p_^2^ =.60; RT: *F*(1,48)=363.90, *p*< .001, η_p_^2^ =.88). RTs but not error rates showed improvements across experimental phases (errors: *F*(1,48)=1.01, *p*= .32, η_p_^2^ =.02; RT: *F*(1,48)=113.49, *p*< .001, η_p_^2^ =.70).

For Demand Avoiders, there was a significant effect of effort level on choice behavior, *F*(5,125)=8.24, *p* < .001. Similar to the pilot behavioral study, they were significantly less likely to choose an effort level with increasing effort requirement; there was a significant linear effect, *F*(1,25)=31.46, *p* < .001. There was a significant effect of effort level on decision time *F*(5,125)=3.05, *p* = .01. The decision time to select an effort level linearly increased with increasing effort levels, *F*(1,25)=6.34, *p*= .02. For Demand Seekers, there was a significant effect of effort level on choice behavior, *F*(3.22,73.98)=4.67, *p* < .01. They were significantly more likely to choose an effort level with increasing effort requirement, there was a significant linear effect, *F*(1,23)=34.20, *p* < .001. The decision time to choose an effort level was similar (*M*= 1.00 s., *SD*= 0.25) across all effort levels, *F*(2.88,66.25)=2.55, *p*= .07.

In order to test selection rates relative to a common baseline, we recalculated overall probability of selecting the easier task separately for only choices when a higher effort level (effort level 2,3,4,5,6) was paired with the lowest effort level (effort level 1). Consistent with our original grouping procedure, the probability of selecting the easier task across these pairs significantly differed between demand groups, with Demand Avoiders selecting the easiest effort level more than Demand Seekers each time the easiest task was paired with a higher effort level (Demand Avoiders: *M*= .72, *SD*= 0.04; Demand Seekers: *M*= .39, *SD*= 0.04; *F*(1,48)=36.76, *p*<.001, η_p_ =.43). Selection rates across demand groups were similar between pairs (*F*(3.42,164.36)=0.36, *p*=.81, η_p_^2^ =.01), indicating that participants’ preferences for the easiest task were stable regardless of the effort level it was paired with.

In order to examine the effects of time on effort selection behavior, overall probability of selecting the easier task was calculated separately for each of the four blocks of the Test Phase. Probability of selecting the easier task did not change across Test blocks (*F*(2.59,124.22)=0.14, *p*=.92, η_p_^2^ =.003), indicating that participants’ effort selection were consistent across time. Furthermore, the consistency in selection behavior did not change between demand groups (*F*(2.59,124.22)=1.60, *p*=.20, η_p_^2^ =.03).

Additionally, we tested the effects of performance and task-switching probability on effort selection behavior using hierarchical regression analysis. First, Error Rates during task-execution did not predict deck selection for either Group (Demand Avoiders: *M*_β_=-0.28., *SD*_β_=0.19, t(156) = −1.48, p = .14; Demand Seekers: *M*_β_=0.08., *SD*_β_=0.11, t(144) = 0.67, p = .50). RT during task execution predicted demand avoidance only in the Demand Avoider group (Demand Avoiders: *M*_β_=-0.44., *SD*_β_=0.09, t(156) = −2.34, p = .02; Demand Seekers: *M*_β_=0.35., *SD*_β_=0.20, t(144) = 1.73, p = .09), indicating that Demand Avoiders selected those tasks that yielded the least time-on-task.

Next, we tested the effects of task-switching probability on the selection rates in the presence of RT in order to see if our linear effort manipulation predicted demand avoidance over and beyond RT during task-execution. Demand Avoiders reliably selected those tasks that required the least task-switching, whereas Demand Seekers sought them (Demand Avoiders: *M*_β_=-0.34., *SD*_β_=0.03, t(156) = −5.82, p < .001; Demand Seekers: *M*_β_=0.24., *SD*_β_=0.06, t(144) = 4.09, p < .001). The effects of RT were not longer significant when task-switching probability was added to the model (Demand Avoiders: *M*_β_=-0.07., *SD*_β_=0.18, t(156) = -0.37, p = .71; Demand Seekers: *M*_β_= 0, *SD*_β_=1.05, t(144) = 0, p = .99), indicating that selection behavior was explained better with our linear task manipulation than time-on-task during task execution.

In summary, the behavioral data provide evidence for two groups of participants that differed in their effort choices despite no demographic or performance differences between them except overall faster RTs for Demand Avoiders. Demand Avoiders (Fig 3B) showed a decreasing tendency to choose a task as more cognitive effort is required. Demand Seekers (Fig 3C) exhibited the opposite pattern (see Supplementary Results 2 for additional analysis). This pattern in effort selections was mostly captured by our linear effort manipulation on task-switching probability for both demand groups. We note that based on a behavioral experiment reported in the Supplementary materials (see Supplementary Results 9), this higher tendency to seek demand in the fMRI group versus the behavioral group may reflect a self-selection bias. Specifically, those who volunteer for fMRI tended to score higher on the Need for Cognition scale, and so might be more likely to be demand seeking. These behaviorally defined groupings were maintained throughout the subsequent fMRI analyses in order to address individual differences.

### 3.2. The functional form of univariate brain activity over effort levels

We investigated the pattern of activation in independent ROIs based on our *a priori* hypotheses, in order to examine their underlying neural function across the execution of increasing effort levels (The results of the selection epoch analysis are reported in Supplementary Results 4). We defined ROIs in the FPN (Network12; Yeo et al, 2011), DMN (Network16; Yeo et al., 2011), and/or ventral striatum (VS; Badre et al., 2014), as described in 2.5.3. Then, using the “Independent Effort Level GLM” (see 2.5.2), we estimated the activation in the network for each effort level separately. These estimates are plotted in Fig 4.

**Figure 4.**
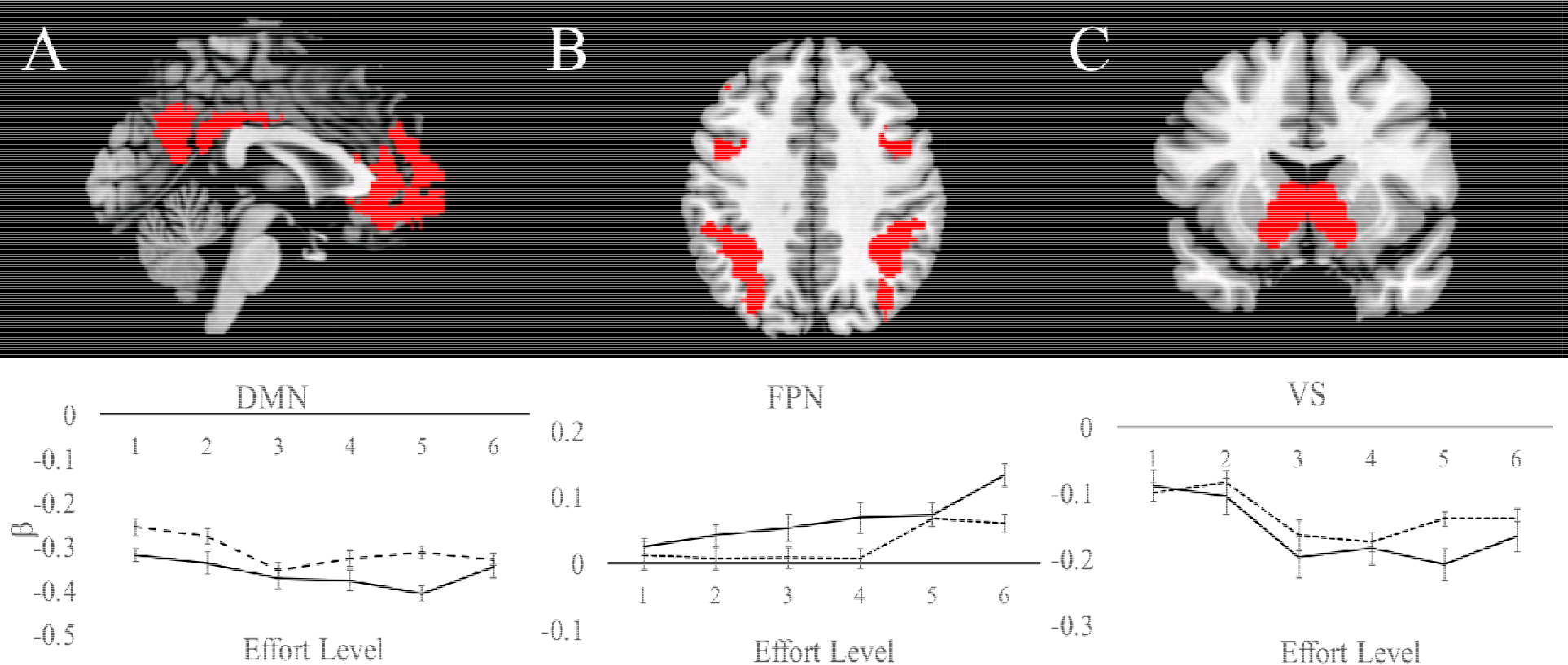
Estimated beta weights at each effort level for both demand groups from independent ROIs defined in (A) the default mode network (DMN), (B) the frontoparietal network (FPN), and (C) the ventral striatum (VS). Demand Avoiders are indicated in solid lines. Demand Seekers are indicated in dashed lines. There was no effect of avoidance group on β estimates for any of the ROIs. In all cases a linear function was the curve of best fit relative to alternatives (summarized in Supplementary Table S3), with the exception of VS. All linear trends are significant at *p* < .05 for both groups.

As expected, the FPN ROI showed a linear trend across effort levels (Fig 4B). There was a significant effect of effort on FPN β estimates, *F*(5,240)=5.19, *p*< .001, η_p_^2^ =.10. The β estimates linearly increased with increasing effort requirements of the task, *F*(1,48)=20.85, *p*< .001, η_p_^2^ =.30. There was no effect of avoidance group on β estimates, *F*(1,48)=0.57, *p*= .45, η_p_^2^ =.01.

The DMN showed a negative linear trend in activation across effort levels (Fig 4A). There was a significant effect of effort on DMN β estimates, *F*(5,240)=4.18, *p*= .001, η_p_^2^ =.08. The β estimates linearly decreased with increasing effort requirements of the task, *F*(1,48)=9.77, p < .01, η_p_^2^ =.17. There was no effect of avoidance group on β estimates, *F*(1,48)=1.06, *p*= .31, η_p_^2^ =.02.

An ROI in VS also showed a negative linear trend (Fig 4C). There was a significant effect of effort on VS β estimates, *F*(3.26,156.52)=5.94, *p* = .001, η_p_^2^ =.11. The β estimates linearly decreased with increasing effort requirements of the task, *F*(1,48)=9.40, *p* < 01, η_p_^2^ =.16. There was no effect of avoidance group on β estimates, *F*(1,48)=0.32, *p*= .58, η_p_^2^ =.01. We note that in all cases a linear function was the curve of best fit relative to alternatives (summarized in Supplementary Table S3), with the exception of VS that, though showing a reliable linear function, was better fit by a quadratic due to an apparent asymptote at level 4.

In order to ensure that our ROI approach did not exclude regions or networks outside of our *a priori* defined ROIs, we conducted whole brain analyses based on parametric contrasts across effort levels. As described in the Supplementary Results 3 and plotted in Figure 5, the results of these whole-brain analysis results were consistent with the ROI analyses.

**Figure 5.**
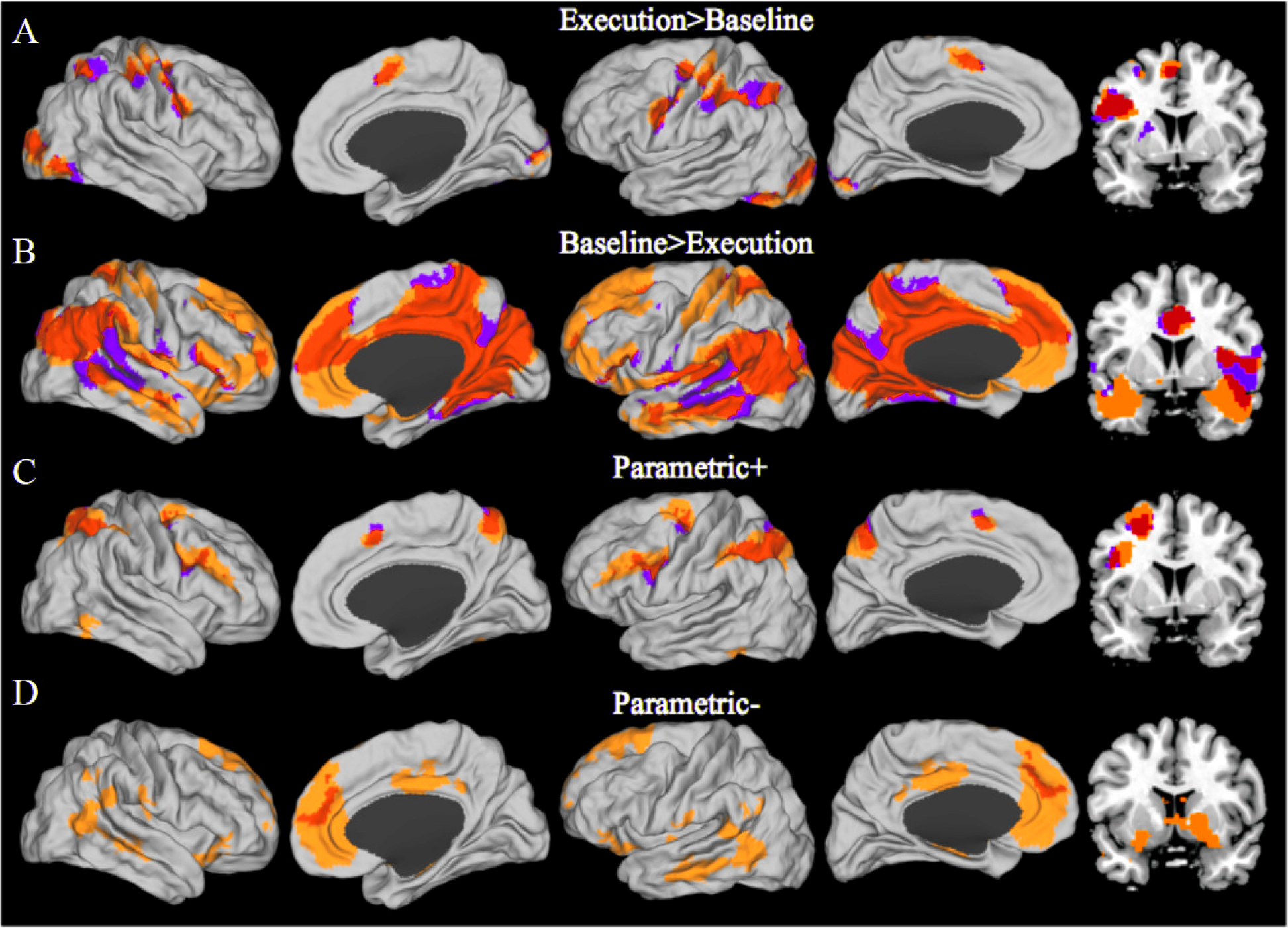
Four whole brain contrasts are plotted on a canonical surface with a coronal slice (Y=8) on the far right to show activation in the basal ganglia. Yellow: Demand Avoiders; purple: Demand Seekers; Red: overlap between Demand Avoiders and Demand Seekers. (A) The contrast of all blocks (across effort levels) from the execution phase versus baseline primarily shows the FPN and other typical task active areas. (B) The reverse contrast of baseline > execution phase blocks shows typical task negative activations including the DMN. (C) The positive parametric effort level effect shows regions that demonstrated a linear increase with effort level, including regions of the FPN. (D) The negative parametric effort level effect shows regions that were negatively correlated with effort level including regions of the DMN and ventral striatum. Images are thresholded at *p* < .05 FWE cluster corrected.

Finally, we conducted a whole brain PCA analysis to ensure that other networks or clusters of voxels do not hold non-linear functions with increasing effort execution that might influence avoidance behavior, but that would not be detected by the linear parametric regressor used in the GLM or by our ROIs. Thus, we adopted a PCA approach to the whole brain analysis. This analysis is used to identify a small set of variables that explain the maximum amount of variance in a large data set. In our case, we used the PCA to identify voxels in the whole brain that share a similar underlying activation functional form across the execution of effort levels.

From 126,866 voxels included in the analysis, the PCA located 3 PCs that explained 90% of the variance of the data. The first PC explained 57% of the variance. The percent variance of the data explained by each subsequent PCs reached asymptote by 4-5 PC, and 5 PCs explained 100% of the total variance (Fig S2). We extracted 3000 voxels that loaded the most positively (Positive PCs) and most negatively (Negative PCs) on each component and extracted their βs in order to observe the functional form of each PC.

Positive PC1 (Fig S3, Table S4) overlapped with the frontoparietal network, and included the left SMA, left superior parietal lobule, left middle frontal gyrus, left inferior frontal gyrus. As we observed in Parametric+ whole-brain analysis, there was a significant effect of effort on FPN cluster β estimates as defined by Positive PC1, *F*(3.84,184.18)=19.08, *p*< .001, η_p_^2^ =.28. The β estimates from these voxels linearly increased with increasing effort requirements of the task, *F*(1,48)=69.24, *p*< .001, η_p_^2^ =.59. This converges with the preceding *a priori* ROI analyses and indicates that the most voxel-wise variance in effort execution can be explained by a linear trend in the brain regions that contribute to controlled behavior.

Negative PC1 (Fig S3, Table S5) overlapped with the DMN, and included the medial prefrontal cortex, posterior cingulate cortex, left and right angular gyrus, left and right temporal gyrus, and left and right inferior frontal gryus. The β estimates linearly decreased with increasing effort requirements of the task, *F*(1,48)=27.45, *p*< .001, η_p_^2^ =.36.

Overall, PCA results indicate that a linear function explained more than 50% of the voxel-wise variance in the shape of activation function across effort levels with Positive PC1 showing a positive trend and Negative PC1 showing a negative trend (see Supplementary Results 5 for Positive and Negative PC2 and PC3). We note that the Positive PC1 has a close correspondence to the *a priori* defined FPN, and the Negative PC1 has a similar relationship to the DMN (see Figure S4). However, these correspondences are not perfect with substantial components of each of the *a priori* networks not showing clear positive or negative linear patterns. The PCA is an unbiased means of defining regions that show a linear pattern across effort levels. Thus, we use the PCA definition for subsequent analyses. However, in order to make its relationship to the *a priori* networks clearer, we refer to these ROI definitions from the PCA as the “Fronto-Parietal Component” and the “Default Mode Component”. We also performed all analyses in the *a priori* defined networks and report these analyses in Supplementary Results 8.

### 3.3. Brain − Behavior Analysis

Having established the functional form of activity change across effort levels, we tested whether brain regions tracking effort execution predicted effort selections. Two correlation techniques were adopted to examine the type of brain-behavior relationship: 1) relationship between brain activity and effort selection rate at each effort level (’Individual task selection’), 2) relationship between change in brain activity and change in effort selection rates across effort levels (‘Effort Level’). As ‘Individual Task Selection’ analysis did not find statistically reliable effects, we report the results of this analysis in Supplementary Results 7.

‘Effort Level’ analysis showed that change in Fronto-Parietal Component during task execution across effort levels (i.e., due to the task switching manipulation) did not reliably predict change in effort selection rates in demand avoiders (r(24) = .17, p=.41). For demand seekers, there was a marginal negative correlation between demand avoidance and Frontoparietal Component change across effort levels (r(22) = -.40, p=.052). Thus, to summarize, we did not find evidence that Fronto-Parietal Component activation and task selection was related to the parametric effort manipulation in demand avoiders. By contrast, demand seeking behavior in demand seekers marginally increased with increasing Fronto-Parietal Component activity change (Fig 6). We emphasize that the latter observation was not statistically significant, and so does not provide evidence of a relationship between activation in FPN and effort selection. Nevertheless, even if one were to accept the marginal effect, the non-significant trend is incongruent with the ‘cost of control’ hypothesis, such that greater FPN recruitment during harder tasks yielded greater effort selection in demand seekers. However, the Demand Group x Fronto-Parietal Component slope interaction was statistically unreliable (*b*= 0.60, *t*(46) = 1.75, *p* = .09), indicating that the relationship between Fronto-Parietal Component slope and demand avoidance did not change as a function of demand group.

**Figure 6.**
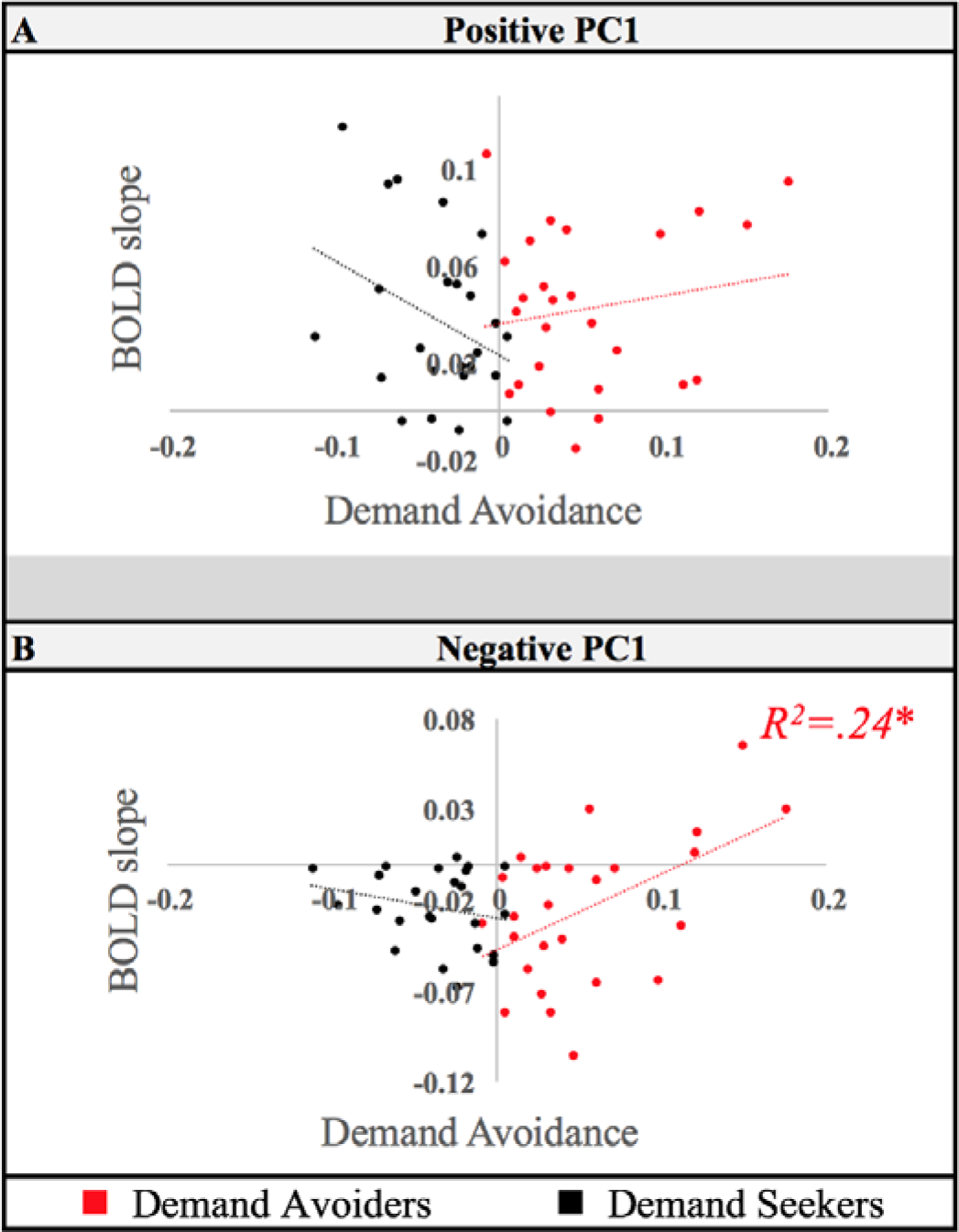
Principal Component cluster correlations with demand avoidance behavior are plotted for (A) Positive (“Fronto-Parietal Component”) and (B) Negative PC1 (“Default Mode Component”). Results for Demand Avoiders are plotted in red. Results for Demand Seekers are plotted in black. The ‘Effort level’ analysis tested whether the change in effort selection rates was correlated to the change of brain activity in each PCA ROI during task execution across effort levels. Default Mode Component showed a positive relationship between the slope of change in this network across effort levels and effort selections. This was only the case in the Demand Avoider group. Error bars plot standard error of the mean. * *p* < .05.

**Table 1.** reports demand avoidance rates for each demand group across runs during the Test Phase.

| Run | Demand Seekers |  | Demand Avoiders |  |
| --- | --- | --- | --- | --- |
|  | *Mean* | *Std. dev.* | *Mean* | *Std. dev.* |
| 1 | 0.44 | 0.12 | 0.59 | 0.11 |
| 2 | 0.42 | 0.10 | 0.60 | 0.12 |
| 3 | 0.41 | 0.09 | 0.63 | 0.13 |
| 4 | 0.44 | 0.07 | 0.59 | 0.17 |

**Table 2.** Whole brain analysis for both demand groups. (* indicates that higher clustering threshold was used in order to break down big clusters for reporting purposes)

| Brain Region | Cluster size | MNI coordinates |  |  | Voxel t-value |
| --- | --- | --- | --- | --- | --- |
|  |  | X | Y | Z |  |
| Execution > Baseline |  |  |  |  |  |
| Demand Avoidants |  |  |  |  |  |
| right occipital cortex | 2320 | 26 | -94 | 2 | 15.78 |
| left occipital cortex | 1502 | -26 | -94 | 2 | 11.74 |
| left supramarginal gyrus | 3830 | -44 | -36 | 42 | 7.19 |
| right superior parietal lobule | 292 | 30 | -58 | 44 | 5.33 |
| Demand Seekers |  |  |  |  |  |
| left occipital cortex | 2998 | -16 | -96 | 0 | 13.21 |
| right occipital cortex | 3502 | -28 | -52 | 46 | 10.57 |
| right superior parietal lobule | 406 | 34 | -56 | 54 | 8.6 |
| left putamen | 373 | -22 | 4 | 10 | 5.6 |
| supplementary motor cortex | 232 | -8 | 10 | 52 | 5.14 |
| Baseline > Execution* |  |  |  |  |  |
| Demand Avoidants |  |  |  |  |  |
| cuneus | 6380 | 16 | -66 | 22 | 14.82 |
| middle cingulate gyrus | 2827 | 0 | -18 | 42 | 13.67 |
| anterior cingulate gyrus | 1856 | -10 | 40 | 14 | 11.97 |
| left angular gyrus | 472 | -44 | -74 | 34 | 11.4 |
| right middle occipital gyrus | 150 | 50 | -70 | 20 | 10.76 |
| left superior frontal gyrus | 316 | -20 | 48 | 34 | 10.09 |
| Demand Seekers |  |  |  |  |  |
| posterior cingulate cortex | 9135 | -2 | -32 | 46 | 17.83 |
| anterior cingulate gyrus | 474 | 2 | 44 | 16 | 12.83 |
| right posterior insula | 731 | 44 | -8 | -4 | 12.47 |
| left supramarginal gyrus | 142 | -60 | -50 | 26 | 12.41 |
| right middle temporal gyrus | 179 | 66 | -20 | -14 | 12.03 |
| right angular gyrus | 381 | 50 | -68 | 28 | 11.47 |
| left middle temporal gyrus | 145 | -58 | -52 | -2 | 11.33 |
| left middle occipital gyrus | 148 | -42 | -78 | 6 | 10.42 |
| Parametric+ |  |  |  |  |  |
| Demand Avoidants |  |  |  |  |  |
| left superior parietal lobule | 3343 | -24 | -64 | 46 | 8.96 |
| left Inferior frontal gyrus | 2146 | -46 | 32 | 18 | 7.73 |
| left supplementary motor cortex | 253 | -8 | 16 | 44 | 5.91 |
| right superior parietal lobule | 697 | 20 | -62 | 54 | 5.62 |
| left inferior temporal gyrus | 288 | -48 | -56 | -14 | 5.3 |
| Demand Seekers |  |  |  |  |  |
| left supramarginal gyrus | 2215 | -40 | -42 | 44 | 7.24 |
| left supplementary motor cortex | 550 | -6 | 14 | 46 | 5.42 |
| left middle Frontal Gyrus | 483 | -42 | 26 | 26 | 5.1 |
| Parametric- |  |  |  |  |  |
| Demand Avoidants |  |  |  |  |  |
| right middle temporal gyrus | 2776 | 52 | -60 | 6 | 7.04 |
| medial superior frontal gyrus | 10580 | 10 | 48 | 16 | 7 |
| right hippocampus | 246 | 32 | -28 | -10 | 6.51 |
| middle cingulate gyrus | 1952 | 4 | -12 | 38 | 6.16 |
| left middle temporal gyrus | 905 | -64 | -20 | -14 | 5.8 |
| left precentral gyrus | 308 | -46 | -10 | 54 | 5.24 |
| left posterior insula | 404 | -38 | -16 | -2 | 4.86 |
| Demand Seekers |  |  |  |  |  |
| left superior frontal gyrus | 456 | -12 | 42 | 26 | 5.13 |

The Default Mode Component, by contrast, did show this relationship (Fig 6). Pearson correlation between Default Mode Component β estimate slopes and demand avoidance behavior across Demand Avoider participants was significant (r(24) = .49, *p* =.01). In other words, those individuals who had shallower Default Mode Component slopes (i.e., more positive, less deactivated) across effort levels during task execution showed greater demand avoidance. This relationship was absent for Demand Seekers (r(22) = -.28, *p* =.19). The Demand Group* Default Mode Component slope interaction was statistically reliable (*b*= 0.99, *t*(46) = 2.44, *p* = .02). In follow up tests, however, we did not find that these differences between the Fronto-Parietal Component and Default Mode Component ROIs were statistically reliable. Thus, we find evidence for a relationship between effort selection and the change in activation across effort levels in Default Mode Component in Demand Avoiders. However, though we do not find evidence of this relationship for Fronto-Parietal Component, we also do not have evidence that this difference distinguishes these brain networks.

Additionally, we tested for the relationship between connectivity within and between Fronto-Parietal Component and Default Mode Component and demand avoidance, in order to see if individual differences in connectivity predict effort selections. Neither connectivity within nor between these networks correlated with behavioral avoidance rates (see Table S6). Thus, together with our analysis on connectivity change across effort levels (see Supplementary Results 6), we found no evidence that changes in connectivity were related to either the experience or behavior related to cognitive effort.

Effortful task-execution naturally comes with performance costs such as increased error rates and response times. Thus, the higher likelihood of time on task or errors might lead to demand avoidance as well. In order to differentiate the role of performance measures from that of Default Mode Component in demand avoidance, we next regressed out the role of performance from brain-behavior relations. A stepwise multiple regression was conducted to evaluate whether Default Mode Component slope, RT slope and ER slope were necessary to predict selection slope in Demand Avoiders. At step 1, the Default Mode Component slope entered into the regression equation and was significantly related to selection slope (Default Mode Component slope: *b* = .59, *t*(24) = 2.78, *p* = .01). The multiple correlation coefficient was .49, indicating approximately 24% of the variance of the selection slope could be accounted for by Default Mode Component slope. At step 2, ER and RT slopes were regressed out of the selection slope, and Default Mode Component slope entered into the regression on the residuals of ER and RT slopes. Default Mode Component slope still reliably predicted the selection slope even after the effects of ER and RT were removed (Default Mode Component slope: *b* = .58, *t*(22) = 2.41, *p* = .03, R^2^=.16), indicating that change in Default Mode Component activity during effort execution predicted change in effort selections over and beyond what could be attributed to the change in performance. However, we do note that the *a priori* ROI definition of DMN no longer predicted demand avoidance when the effects of performance were controlled (*p* = .09). As discussed below, this difference likely arose due to the substantially broader ROI for the *a priori* definition.

Next, we tested the effects of performance and task-switching on Fronto-Parietal Component activity. For Demand Avoiders, task-switching probability predicted Fronto-Parietal Component recruitment over and beyond performance measures (*M*_β_=0.20., *SD*_β_=0.06, t(153.86) = 3.10, p < .01). For Demand Seekers, both ER and task-switching probability explained FrontoParietal Component recruitment (ER: *M*_β_=-0.90, *SD*_β_=0.32, t(15.5) = −2.76, p = .01; TS: *M*_β_=0.24., *SD*_β_=0.07, t(60.43) = 3.51, p < .001). Comparison of the models with ER and without ER showed that, the inclusion of ERs in the model predicting Fronto-Parietal Component significantly improved goodness of fit compared to the model that only included task-switching probability (X^2^ (2) = 12.09, p < .01), indicating that for Demand Seekers, Fronto-Parietal Component recruitment was best explained by a combination of task-switching probability and error signals. Thus, for demand seekers, effort tasks that yielded greater task-switching probability and smaller error rates predicted greater Fronto-Parietal Component recruitment during task-execution. In other words, FPN recruitment increased during blocks participants showed greater accuracy on difficult trials.

Overall, we have observed that change in Default Mode Component predicts change in effort selection as a function of effort levels in Demand Avoiders over and beyond what can be explained by performance measures ER and RT. Fronto-Parietal Component activation did not predict effort selections in Demand Avoiders, and marginally predicted demend seeking behavior in Demand Seekers.

## Discussion

Here, we aimed to rigorously test the ‘cost of control’ hypothesis of effort by using a parametric version of DST and observing brain activity in two separate epochs of effort-based-decision-making. The ‘Cost of control’ hypothesis makes two core proposals: 1) effort registers as disutility by the brain, 2) the cost derives from cognitive control demands. Thus, we predicted that increasing effort would decrease brain activity in the reward network and increase control-related brain activity in FPN. And, further, FPN-activity due to effort execution would predict effort selection. We found only partial support for these predictions, and rather located unexpected evidence that engagement of the DMN (or failure to disengage) due to effort during a task influences effort-based decisions. We consider each of these findings in turn.

Consistent with the idea that effort might register as disutility, we observed that reward network, including VS, vmPFC and PCC, linearly reduced activity as a function of increasing effort execution, providing support for the ‘cost of effort’ hypothesis and consistent with prior reports (Schouppe et al., 2014; Botvinick et al., 2009). We also observed a saturating decrease in VS activation as effort levels increased, consistent with observations by Botvinick et al. (2009). However, VS activity during task execution did not predict effort selections (Supplementary Results 8), indicating that effort costs, as tracked by VS during task-execution, do not directly influence demand avoidance behavior in DST paradigms in the absence of monetary reward. Nevertheless, we only scanned during the “Test phase” when the association of effort with the different task conditions had already been established. It is reasonable to hypothesize that the costs computed by VS may be differentially important during the “Learning phase” when effort associations are being acquired. This will be an important target of future research.

To test the hypothesis that effort costs arise from engagement of the cognitive control system, we manipulated effort by varying cognitive control demands in a task-switching paradigm. Consistent with the literature that implicates FPN activity in tasks demanding cognitive control, we observed that most of the variance in our data could be explained by a linearly increasing activation function in this network with increasing task-switching probability.

Under the cost hypothesis, we would expect that effort seekers would be different from Demand Avoiders in four ways: less FPN activity, smaller error rates, and shorter response times or greater VS activity. These neural and behavioral markers would indicate that Demand Seekers either exerted less effort than Demand Avoiders or that they valued effort exertion more than Demand Avoiders. However, the results show that there was no difference in error rates or neural recruitment between demand groups. Moreover, contrary to the expectations of the cost hypothesis, Demand Avoiding participants had overall faster response times compared to Demand Seeking participants, and if anything, Demand Seekers who showed greater demand seeking behavior showed an increasing trend in FPN recruitment across the execution of effort levels. These results suggest that behavioral and neural proxies for cognitive control recruitment may not obligatorily register as something costly. Instead, difficult tasks that yield greater change in FPN recruitment might be preferred by demand seeking individuals.

Further, the linear increase in FPN activity due to the cognitive control manipulation was not predictive of an increase in demand avoidance behavior. And, we found that for Demand Seekers, performance-related factors like error likelihood predicted FPN activity even after the effects of our experimental cognitive control manipulation were controlled. Thus, we observe only partial support for the cost of control hypothesis, at least as it might stem from engagement of the FPN network. Performance indices like error-likelihood and RT likely correlate with cognitive control demands in a task, but they could relate to other factors, as well. Thus, to avoid circularity in relating cognitive control to FPN activation and demand avoidance, the task switching manipulation provided an independent definition of cognitive control. And using this definition of cognitive control demands, we did not find evidence that changes in activation in FPN attributable to cognitive control were related to demand avoidance.

We note that our observations are not necessarily inconsistent with prior reports. A previous study that also utilized a task-switching paradigm to manipulate cognitive effort (McGuire and Botvinick, 2010) showed that a change in FPN activity predicted demand avoidance. However, this study used a binary comparison, where a change in brain activity and task selections between low and high effort conditions was not corrected by the change in intermediate task levels. By contrast, our parametric manipulation provided greater sensitivity to separate changes in FPN activation due to the cognitive control manipulation versus other factors related to performance.

Additionally, our study is the first to test the neural mechanisms underlying effort selection and effort execution in the same experiment. Whole-brain analysis of the Selection Epoch showed that execution and the selection of effort did not yield activity in the same brain regions. While the execution of effort yielded parametric effects in FPN, DMN and VS, the selection of effort yielded parametric effects only in the primary motor cortex, suggesting that the selection and the execution of cognitive effort recruit different neural systems. These differences might help explain the observed discrepancies in the literature regarding differential the engagement of brain networks depending on whether one is performing or anticipating an effortful task. Future studies should attend to temporal dynamics of effort performance.

There are no previous studies that parametrically manipulated implicit cognitive effort demands in the absence of monetary reward as we did here. However, some studies have manipulated explicit cognitive effort demands using reward discounting procedures (Chong et al., 2015; Apps et al., 2015; Massar et al., 2015; Libedinsky et al., 2013; Westbrook et al., 2013). For example, Cognitive Effort Discounting (COG-ED) paradigm (Westbrook et al., 2013) shows that participants parametrically discount task value by increasing demands on cognitive effort. This is consistent with our finding that probability of selecting a task parametrically decreases for demand avoiding participants. However, unlike the discounting procedures, we also show that the same task demands were found increasingly valuable by demand seeking participants. This individual difference suggests that effort is not obligatorily aversive or coded as disutility, but rather effort signals can be interpreted positively or negatively depending on the participants.

The discrepancies in these observations might be due to three notable differences between this procedure and the one employed in discounting paradigms, beyond our inclusion of fMRI participants. First, in COG-ED, participants were explicitly aware of the difficulty difference between levels, as the instructions regarding each effort level significantly differed due to the nature of the effort task. In our version of DST, the task instructions were the same across effort levels, and as post-experimental debriefing inventories showed most participants were not explicitly aware of a difficulty difference between effort levels. Second, in COG-ED, participants made their choices without immediately executing the chosen option, while our procedure required the participants to execute the effort level immediately following their selections Third, in COG-ED, participants made a choice between an option that yields low effort and low reward, and an option that yields large effort and large reward. Our task was performed in the absence of explicit feedback or reward. Directly testing these dimensions of decision making may be fruitful directions for future research.

Further, a notable advantage of the effort discounting procedures such as COD-ED is that they can estimate the subjective point of equality for each effort level by requiring choices between options that yields low effort and low reward, and an option that yield large effort and large reward. However, in doing so, COG-ED procedure must assume that the inclusion of extrinsic reward does not influence the intrinsic evaluation of cognitive effort. The advantage of the current task is that it aims to test the intrinsic cost/value associated with effort in the absence of feedback or incentives, and thus is directly testing the influence of intrinsic motivation that would drive voluntary effort selections in the absence of monetary reward. Nevertheless, without a discounting approach, we are unable to assess the unit of exchange between effort and monetary reward across effort levels.

A related open question concerns whether demands on cognitive control are themselves effortful, or rather, cognitive control tends to consume more time and it is the time on task that the brain taxes as costly. Kool et al. (2010) showed that participants avoided switching between tasks even when staying on task meant longer task durations. Consistent with those findings, we observed that the effect of task-switching probability predicted effort selections, however in opposite directions for each demand group. While Demand Avoiders avoided those tasks that yielded greater task-switching, Demand Seekers chose them. Future studies should aim at equating error-rates and time-on-task across effort levels in order to empirically differentiate the effects of task performance from cognitive control recruitment on effort decisions.

Whereas we found partial support for the cost of control hypothesis with regard to engagement of the FPN, a novel discovery in the present study was that the DMN is a robust correlate of effort avoidance. Specifically, demand avoiding participants who showed a diminished negative change in DMN activity across cognitive control defined effort levels showed the highest demand avoidance rate. The relationship between DMN and effort selections persisted even after performance measures such as RT and ER were controlled. We note that though FPN did not show these effects, as discussed above, we also did not find a significant difference between these networks. So, we cannot conclude that this is a qualitative difference such that DMN is more determinant of effort selection than FPN. These potential differences may be an important avenue for future research.

It is not evident from our study what drives this relationship between DMN activity, though prior studies of this network may offer some clues. While FPN recruitment has been shown to underlie cognitive control, DMN has been associated with a wide range of internal mental functions, including self-referential thoughts, mind-wandering, and episodic future planning (Buckner et al., 2008; Weismann et al., 2006). Inhibition of DMN has been argued to support task-related attention (Spreng, 2012), while the inability to inhibit DMN has been related to dysfunctional cognitive processes such as rumination (Nejad et al., 2013) and pathologies such as depression (Anticevic et al., 2012; Lemogne et al., 2012). Accordingly, in our study, those participants who showed higher effort avoidance rates could have had increased difficulty deactivating DMN activity or relied more on the processes it entails, which in turn might have registered as a cost. Future studies should seek to replicate this discovery and to determine what factor drives this change in DMN activity across effort levels.

It is possible that this DMN activity is related to reward and/or value processing that predicts effort selections. A common observation in the literature is that vmPFC positively tracks subjective value and predicts subjects’ preferences between choices (Rushworth et al., 2011). Given this functional association, the ‘cost of effort’ hypothesis would predict that individuals will avoid tasks that yield the least activity in the reward network. However, in our study, we observed that individuals who showed the least reduction in this network showed the greatest demand avoidance. In addition to linear reductions in vmPFC and PCC activity with increasing effort, independent ROI definitions of VS showed that VS reduces its activity across increasing effort, however, DMN but not VS predicted demand avoidance behavior (see Supplementary Results 8). These results together suggest that while reward network tracks effort in congruence with the ‘cost of effort’ hypothesis, this cost does not predict effort selections.

We also probed the relationship of functional connectivity to effort avoidance. Stronger functional or structural connectivity within-FPN has been associated with working memory performance and general intelligence (Cole et al., 2012; Gordon et al., 2012; Nagel et al., 2011), executive capacity (Gordon, Lee, et al., 2011), cognitive dysfunctionalities (Rosenberg et al., 2016) and procrastination behavior (Wu et al., 2016). However, we did not find evidence that connectivity within the FPN network or between FPN and DMN networks predicted effort selections.

Finally, individual differences are an important variable considered here that has not received much attention in the literature. In our fMRI task, we observed high individual variability in effort avoidance, such that roughly half the participants were better characterized as Demand Seekers than Demand Avoiders. This rate of variability has not been reported previously in the literature and was not the case in our own behavior-only task (Experiment 1).

The conflicting results might be due to at least three reasons: 1) self-selection bias in fMRI experiments, 2) context effects in fMRI settings, 3) time-of-year effects for subject pool participants who showed a tendency to participate in our experiments around their finals. The behavioral participants in our behavior-only study mostly consisted of undergraduate participants who volunteered for course credit. However, our fMRI participants consisted of a more variable sample who volunteered for fMRI scanning at different months of the year in response to ads placed around campus. A recent study showed that PET scan volunteers significantly scored higher in sensation-seeking trait compared to behavior-only volunteers (Oswald et al., 2013). Sensation-seeking is also a trait that positively correlated with Need for Cognition (NfC), a selfreport inventory that tests effort avoidance and that negatively correlates with Effort Discounting scores (Schuller, 1999).

We confirmed in a follow up study that an independently recruited sample of volunteers responding to advertisements for fMRI scored significantly higher on NfC compared to subject pool participants (see Supplementary Results 9). Of course, this study does not provide direct evidence that the fMRI participants reported in this paper had higher NfC, as we did not administer this to them at the time they were scanned. Thus, we can only provide the general observation that NfC is higher among participants responding to ads for fMRI. Future studies should directly test the relationship between the demand avoidance scores in this parametric version of DST and NfC.

The demonstration of individual differences is central to the current study for three reasons. First, we think that individual differences in effort expenditure behavior and brain activity has not been fully explored in the effort literature, and ours is the first study to show that the same performance and brain activity utilized by Demand Avoiders to avoid effort, can be used by Demand Seekers to choose effort. Second, for the half of our sample that sought effort, increased frequency of task-switching and FPN recruitment registered as something valuable rather than costly. This observation was counter to expectations and strains against strong versions of the ‘cost of effort’ hypothesis. Third, the results show that the DMN-behavior relationship we observe in Demand Avoiders is not present in Demand Seekers, which indicates that DMN-based activity can register as an exclusive cost signal that is tracked by demand avoiding individuals.

Regardless of the source of the individual differences, we were able to test how the factors affecting effort-based decisions differed across these groups. Although there were no differences in brain activity and error rates between demand groups during task execution, Demand Avoiders avoided those tasks that yielded greater task-switching, while Demand Seekers chose them. This discrepancy in the way separate demand groups utilized the same information indicates that FPN, and task performance could influence effort-based value-based decisions generally, even if not necessarily in terms of a cost signal. On the other hand, reduced DMN de-activation across effort levels influenced effort avoidance only in demand avoiding participants, indicating that change in DMN activity across effort levels entered into effort-based decisions exclusively as a cost signal. However, note that the findings of our PCA-based ROI definition of DMN and a priori definition of DMN are not completely congruent. While both definitions showed that those participants who showed less DMN inhibition across effort levels showed greater demand avoidance rates, this relationship persisted only for the PCA-defined DMN when the effects of performance were controlled. The effect was only marginal using the a priori DMN definition. This discrepancy is likely due to differences in the brain regions these ROIs encompass. The a priori DMN definition covers a broader region than that encompassed by the PCA regions (Figure S4).

A potential caveat given that our effort task parametrically manipulated implicit effort costs, is that a participant with consistent but arbitrary rankings might show a linear slope across effort selection while they also showed a bias towards or against selecting the easier task. One source of such stable preferences unrelated to effort could stem from a preference for the symbol shapes that cued effort levels. However, symbol orders were randomized across participants, making it unlikely that a group of participants would systematically prefer symbol shapes that also correlated with our effort manipulation. Further, these preferences would also evidently correlate with changes in brain during execution that we report here. Thus, effort avoidance provides a parsimonious account of the full set of results we present here. Nevertheless, we do note that it is possible that effort selections could have been driven by a fixed, but arbitrary, preference ranking.

In conclusion, we adopted a parametric version of DST in order to test the ‘cost of control’ hypothesis and explore the neural mechanisms that underlie effort execution. We have observed that the reward network reduces activity in response to executing more effortful tasks, in congruence with the ‘cost of effort’ hypothesis. DMN but not FPN predicted effort avoidance behavior in demand avoiding participants, indicating that control recruitment as indexed by FPN recruitment does not underlie demand avoidance. As behavioral task performance can represent the required cognitive control to perform well at a task, increased time-on-task and error-likelihood could constitute costs associated with the opportunity-cost of time and error-avoidance. However, neither error rates or response times predicted effort selections in either Demand Avoiders or Demand Seekers. Additionally, high individual variability in effort avoidance behavior in our task showed that the direction greater task-switching probability influenced effort-based decisions depended on the demand group, indicating that proxies of cognitive control do not exclusively influence effort-based decisions as a cost. On the other hand, it was shown that reduced DMN de-activation was an exclusive cost signal that differentiated Demand Avoiders from Demand Seekers, promising to be a new avenue for future effort-based research.

## Acknowledgements

We are grateful to members of the Badre lab, the Frank lab, and the Shenhav lab for their helpful interactions over this work. In particular, we thank Amitai Shehav and Michael Frank for their insightful feedback. We also thank Britanny Ciullo and Nada Hamzah for their assistance with data collection. The present work was supported by R01 awards from the National Institute of Neurological Disease and Stroke (NS065046) and the National Institute of Mental Health (MH099078), a MURI award from the Office of Naval Research (N00014-16-1-2832), and a fellowship from the James S. McDonnell Foundation. CS was also supported through a T32 institutional training grant from the NIH (3T32NS062443-08S1).

